# Effect of Biologically Active Hydrogen-Generating Powder Mixed with Food in a Model of Liver Injury in Rats

**DOI:** 10.1101/2021.04.07.438869

**Authors:** Bronislava Gedulin, Marina Safonov, Vladimir L. Safonov

**Affiliations:** Satiogen Pharmaceutical Inc, San Diego, California, USA; H2 Universe LLC, Granbury, Texas, USA

**Keywords:** Molecular hydrogen, magnesium, liver injury, biomarkers

## Abstract

**Background:** Hydrogen was determined to have good efficacy for reducing key blood level biomarkers associated with liver injury suggesting the compound may provide novel option for the treatment of several liver diseases by decreasing of accumulation of toxins and reduction of levels of serum liver enzymes.

**Materials and methods:** The present pharmacological study was conducted to evaluate the effectiveness of a gastric hydrogen generating powder called “AquaActive” (AA) in a rat model of liver injury, the partial bile duct ligation (pBDL), following administration of 0.2% AA formulated in rat pellets food for 14 days.

**Results:** The data indicate that treatment with AA at a dose of 150 mg/kg/day can effectively slow the progression of liver injury that is triggered by bile duct ligation in rats. At both 7 and 14 days post-pBDL surgery, treatment with AA exhibited reductions in ALT, AST, ALP, GGT and total bilirubin, most of which were statistically significant. At 7 days, the compound showed statistically significant decreases in ALP, GGT and total bilirubin levels. Although the values of some parameters decreased in the vehicle group by 14 days, additional reductions due to AA treatment were sustained for ALP, AST and for GGT and total bilirubin. GGT and total bilirubin level after 14 days of treatment compared to the vehicle-treated control group were observed to be highly significant (p<0.05).

**Conclusion:** Thus AA, gastric hydrogen generating powder demonstrated a good efficacy for reducing key parameters associated with liver function in the pBDL model.

## 1. Introduction

For a long time, molecular hydrogen (H_2_) was assumed to be inactive and useless in the cells and organs of mammals. However, recent studies reversed this notion by representing that H_2_ reacts with extremely reactive oxygen species (ROS) like hydroxyl radical (•OH) and peroxynitrite (ONOO-) in cells. Basic and clinical research has revealed that molecular hydrogen demonstrates important medicinal properties with antioxidant, anti-inflammatory and anti-apoptopic protective effect on cells and organs [1-5]. To date, about two thousand scientific articles on the direction of hydrogen therapy have already been published on experimental animal models and human studies.

The liver is one of the vital organs that can be subject to many diseases and injuries. In this paper, we look at the effects of molecular hydrogen in a rat liver injury model. It should be noted that the therapeutic role of molecular hydrogen has already been reported in several works. Fukuda et al. [6] showed that the inhalation of hydrogen gas is applicable for hepatic injury caused by ischemia/reperfusion, using mice. Zhang et al. [7] demonstrated using rats that hydrogen gas inhalation protects against liver ischemia/reperfusion injury by activating the NF-κB signaling pathway. Liu et al. [8] have studied how hydrogen-rich saline protects against liver injury in rats with obstructive jaundice. Sun et al. [9] showed that hydrogen-rich saline could protect against liver injury in mice and inhibit the processes leading to liver cirrhosis and hepatocyte compensatory proliferation. Tan et al. [10] have shown that hydrogen-rich saline attenuates postoperative liver failure after major hepatectomy in rats. Yu et al. [11] demonstrated that lactulose accelerates liver regeneration in rats by inducing hydrogen. Uto et al. [12] observed that hydrogen-rich solution attenuates cold ischemia-reperfusion injury in rat liver transplantation. In this paper we study model of liver injury in rats adding gastric hydrogen generating powder in their food. We believe this approach provides a simple and controlled method of hydrogen therapy for the liver.

## 2. Models and known treatment

One of the liver’s roles involves removing and breaking down most drugs and chemicals from bloodstream. Breaking down toxins creates chemicals that can damage the liver; one of the manifestations of liver damage is cholestasis and jaundice.

Our acute model liver injury mimics conditions of Toxic hepatitis that can occur when liver develops inflammation because of exposure to a toxic substance, acute complication of hepatitis A, hepatitis B and hepatitis E can cause acute liver failure, metabolic and cholestatic liver diseases (autoimmune etiology) also leads to progressive liver injury.

Our acute model liver injury mimics many conditions of liver injury described below.

### 2.1 Toxic hepatitis

Toxic hepatitis can be caused by:

- Alcohol. Heavy drinking over many years can lead to alcoholic hepatitis - inflammation in the liver due to alcohol.
- Over-the-counter pain relievers. Nonprescription pain relievers such as acetaminophen (Tylenol, others), aspirin, ibuprofen (Advil, Motrin IB, others) and naproxen (Aleve, others) can damage your liver, especially if taken frequently or combined with alcohol.
- Prescription medications. Some medications linked to serious liver injury include the combination drug amoxicillin-clavulanate (Augmentin), halothane, isoniazid, valproic acid (Depakene), phenytoin (Dilantin, Phenytek), azathioprine (Azasan, Imuran), niacin (Niaspan), atorvastatin (Lipitor), lovastatin (Mevacor), pravastatin (Pravachol), simvastatin (Zocor), fluvastatin (Lescol), rosuvastatin (Crestor), ketoconazole, certain antibiotics, certain antivirals and anabolic steroids. There are many others.
- Herbs and supplements. Some herbs considered dangerous to the liver include cascara, chaparral, comfrey, kava and ephedra. There are many others. Children can develop liver damage if they mistake vitamin supplements for candy and take large doses.
- Industrial chemicals. Chemicals people exposed to on the job can cause liver injury. Common chemicals that can cause liver damage include the dry-cleaning solvent carbon tetrachloride, a substance used to make plastics called vinyl chloride, the herbicide paraquat and a group of industrial chemicals called polychlorinated biphenyls
- Metabolic diseases. Rare metabolic diseases, such as Wilson’s disease and acute fatty liver of pregnancy, infrequently cause acute liver failure.

### 2.2 Cholestatic Liver Disease

Impairment of the egress of bile acids from the liver leads to cholestasis, hepatocellular injury and damage, and progressive liver disease that may ultimately lead to the need for liver transplantation. Primary biliary cirrhosis (PBC) and primary sclerosing cholangitis (PSC) represent examples of chronic cholestatic liver disease in adults. Itch is the archetypal symptom of cholestasis, occurring at all stages of cholestatic liver disease, with or without jaundice. Its etiology remains unclear.

### 2.3 Primary Biliary Cirrhosis

Primary biliary cirrhosis (PBC) is a chronic and slowly progressive cholestatic liver disease of autoimmune etiology characterized by injury of the intrahepatic bile ducts that may eventually lead to liver failure. Affected individuals are usually in their fifth to seventh decades of life at time of diagnosis, and 90% are women.

The majority of patients are asymptomatic at diagnosis; however, some patients present with symptoms of fatigue and/or pruritus. Laboratory abnormalities include elevations in bile acids, alkaline phosphatase, bilirubin and AST. UDCA has been shown to improve serum biochemical markers and can slow the disease progression in many but not all patients.

### 2.4 Treatment of liver injury

Known Rx treatment of liver injury is very limited. Bile salt resins and ursodeoxycholic acid (UDCA) are only FDA approved medication for adult treatment of progressive cholestatic liver disease has been shown to improve serum biochemical markers and can slow the disease progression in many but not all patients [13] also UDCA shown cytoprotective, antiapoptotic, and immunomodulatory properties [14]. UDCA usually provides only limited relief of itching, in spite of additional treatment with anti-pruritic and bile salt resins [13].

### 2.5 Cholestasis

Cholestasis is an impairment or cessation of bile flow that causes hepatotoxicity due to accumulation of bile acids and other toxins in the liver. Cholestasis is a detrimental component of many liver diseases, including cholelithiasis, intrahepatic cholestasis of pregnancy, primary biliary cirrhosis (PBC) and primary sclerosing cholangitis (PSC). In addition to significant loss of liver function which can ultimately require a liver transplant, patients with these diseases suffer to various degrees with pruritus (severe itching), jaundice, fat-soluble vitamin deficiency and lethargy.

## 3. Materials and methods

Our study determines inflammatory and protective effect of orally administered gastric hydrogen generating powder called “AquaActive” (AA) administrated with food in the form of pellets in rat model of liver injury induced by extrahepatic cholestasis.

### 3.1 Hydrogen generating powder, AA

AquaActive powder was provided by DIBAL, LLC (San Diego, California, USA). It is one of the patented dietary products [15] that, when dissolved in aqueous solutions give negative redox and release molecular hydrogen. Some of the physicochemical properties of AA were studied in Ref. [16]. AA composition comprises of potassium bicarbonate, sodium bicarbonate, tartaric acid, l-leucine, organic sea salt, calcium lactate, inulin and metallic magnesium particles (40 μm). The latter are responsible for the generation of molecular hydrogen in gastric juice (Mg + 2HCl = MgCl_2_ + H_2_). 200 mg of AA when dissolved in gastric juice (hydrochloric acid) gives about 0.68 mmol of molecular hydrogen.

### 3.2 Animals

All procedures were conducted according to the guidelines of the Institutional Animal Care and Use Committee at Absorption Systems, Inc. (San Diego, California, USA). Male 6-wk old Sprague Dawley (HSD) rats (Harlan, Indianapolis, Indiana, USA) were acclimated to the facility for one week and then fasted overnight before initiating surgical procedures.

### 3.3 Experimental Procedures

Rats were anesthetized with inhalation of isoflurane and maintained under anesthesia throughout the procedure. All surgeries were performed using aseptic techniques including sterile materials, instruments and appropriate personal protective equipment. The rats receive an analgesic dose of ketoprofen (5 mg/kg subcutaneously) immediately following surgery with a second dose repeated approximately 24 hours post-surgery.

### 3.4 Partial bile duct ligation (pBDL) to induce experimental cholestasis

The partial bile duct ligation (pBDL) procedure allows narrowing the bile duct lumen without completely closing it as is the case for a total ligation. The bile duct was exposed via a standard midline laparotomy incision and the common bile duct isolated. A segment of polyethylene tubing (P.E. 10-0.110 mm OD pre-sterilized R-JVC STD, Braintree Scientific) was placed in parallel next to the isolated bile duct. The bile duct and the PE tubing placed adjacent to the vessel were ligated at a single location using 4-0 silk braided suture. After tight fixation of the ligature, the PE tubing was carefully removed resulting in a constriction of the bile duct lumen.

Sham laparotomies were performed, and the common bile duct isolated to simulate the ligation procedure without placing a ligature around the bile duct.

### 3.5 Treatment

After surgery rats were randomly distributed into two groups which were fed on either a normal diet as food pellets (group of 7 rats) or a food formulated with 0.2% AA to the pellets (group of 6 rats). Rats were maintained on this dietary regimen for 14 days and then sacrificed. Sham operated animals (group of 3 rats) were fed with normal non formulated food pellets.

Calculated daily dose of AA blended with pellets was 150 mg/kg/day. Daily rats consumed approximately 25 g of food containing AA. Rat weight is about 320 g (6-8 weeks old rats. All animals were given regular drinking water provided by CRO.

### 3.6 Formulation

200 mg of AA was blended with 100 g of the food pellets and placed in the cage of each treated rat in the small container. Food was added to the container every day in duration of total 14 days of treatment. The groups received water ad libitum. Food consumption was controlled every day.

### 3.7 Samples collection

Blood (0.5 ml) was collected from the saphenous vein on days 0, 7 and 14 after surgery in the 1 ml Eppendorf tubes, samples were centrifuged within 10 minutes at 2,000 x g for 15 minutes at 4° C, serum was immediately separated and stored at −80°C for later analysis. On day 14, after surgery animals were euthanized with an overdose of isoflurane and their livers collected and fixed for future histological analysis.

### 3.8 Biochemical Analysis

Serum samples were prepared from the blood and analyzed at Antech Diagnostic (Irvine, California, USA) for the serum enzymes alanine aminotransferase (ALT), aspartate aminotransferase (AST), alkaline phosphatase (ALP), α-glutamyl transpeptidase (GGT) and for total bilirubin (Total Bil).

### 3.9 Statistical analyses

Results are presented as means ± SEM and analyzed for statistical significance using an unpaired Student’s t-test analysis in GraphPad Prism 5. Statistical differences for the comparison between two groups were attained if P<0.05.

## 4. Results

As shown in Figure 1, by seven days after pBDL surgery, rats exhibited marked elevations of liver function parameters compared to the Sham control group indicative of progressive liver injury due to partial ligation of the bile duct. Serum enzymes ALT, AST were increased by approximately 2-fold, ALP level was increased by 5-fold, GGT was increased by 6-fold and total bilirubin was increased by 50-fold (all parameters had P<0.001, vs. Sham operated animals). Notably, treatment with 0.2% AA blended in rat food after 7 days of treatment showed a trend toward reduction of all parameters, the changes in several parameters had reached statistical significance as ALP, GGT and total bilirubin levels were decreased by 45.6%, 31.9%, and 45.8% in comparison to the vehicle control group of pBDL rats (P=0.008, P=0.016, P=0.003, respectively). (Table 1, Figure 1).

**Table 1.** Liver panel test results 7 days after liver injury and treatment with 0.2% of AA formulated in rat food pellets.

|  | Sham n=3 |  |  | Control Vehicle n=7 |  |  | 0.2% AA powder Feed n=6 |  |  |
| --- | --- | --- | --- | --- | --- | --- | --- | --- | --- |
|  | Mean | sem | n | Mean | sem | n | Mean | sem | n |
| AST | 149.0 | 27.2 | 3.0 | 357.1 | 32.6 | 7.0 | 272.5 | 37.3 | 6.0 |
| ALT | 59.0 | 9.5 | 3.0 | 120.4 | 8.2 | 7.0 | 93.0 | 7.5 | 6.0 |
| ALP | 26.3 | 4.7 | 3.0 | 193.4 | 25.5 | 7.0 | 105.2 | 14.8 | 6.0 |
| GGT | 2.7 | 0.9 | 3.0 | 16.9 | 2.4 | 7.0 | 11.5 | 1.6 | 6.0 |
| Total Bil | 0.2 | 0.1 | 3.0 | 10.7 | 0.2 | 7.0 | 5.8 | 1.6 | 6.0 |

**Figure 1.**
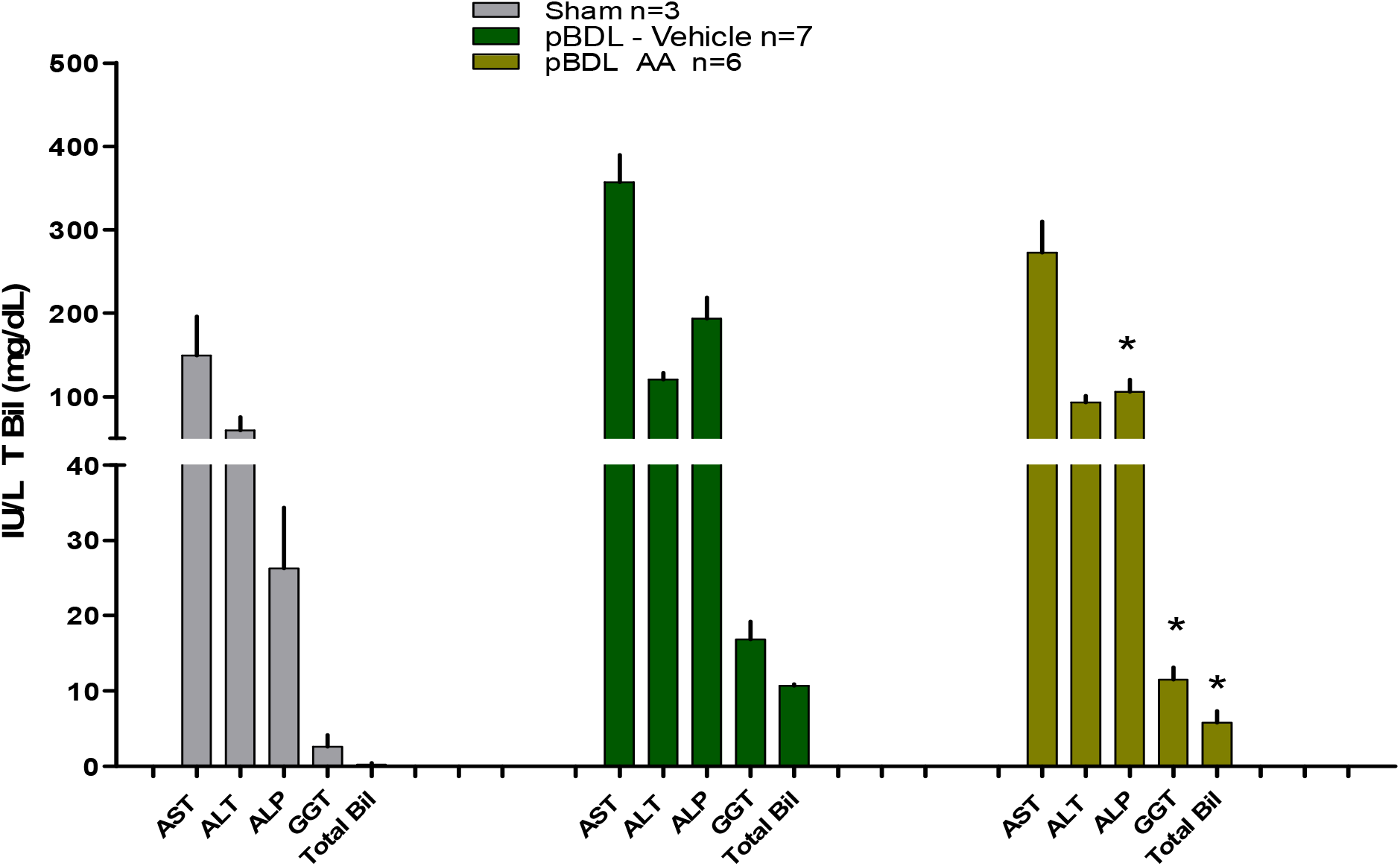
Effect of 0.2% AA formulated in rat food pellets liver tests parameters after 7 days after pBDL surgery. Sham control data is provided to show effect of pBDL on these parameters.

By 14 days post-surgery, animals treated with AA continued to show reductions in all liver function parameters compared to the vehicle-treated control group (Table 2). AST, GGT and total bilirubin levels were decreased by 42.5%, 49.6%, and 64.1% in comparison to the vehicle control group of pBDL (Table 2, Figure 2). GGT and Total bilirubin level both reached statistical significance (P<0.05) compared to the vehicle-treated control group (Figure 2).

**Table 2.** Liver panel test 14 days after liver injury and treatment with 0.2% AA formulated in rat food pellets.

|  | Sham n=3 |  |  | Control Vehicle n=7 |  |  | 0.2% AA Powder Feed n=6 |  |  |
| --- | --- | --- | --- | --- | --- | --- | --- | --- | --- |
|  | mean | sem | n | mean | sem | n | mean | sem | n |
| AST | 62.0 | 3.1 | 3.0 | 261.6 | 32.5 | 7.0 | 150.7 | 32.8 | 6.0 |
| ALT | 55.3 | 2.7 | 3.0 | 80.0 | 4.1 | 7.0 | 65.8 | 5.7 | 6.0 |
| ALP | 258.7 | 25.9 | 3.0 | 443.0 | 33.9 | 7.0 | 382.0 | 35.9 | 6.0 |
| GGT | 1.0 | 0.0 | 3.0 | 22.4 | 3.4 | 7.0 | 11.3 | 4.6 | 6.0 |
| Total Bil | 0.1 | 0.0 | 3.0 | 10.5 | 1.3 | 7.0 | 3.7 | 2.1 | 6.0 |

**Figure 2.**
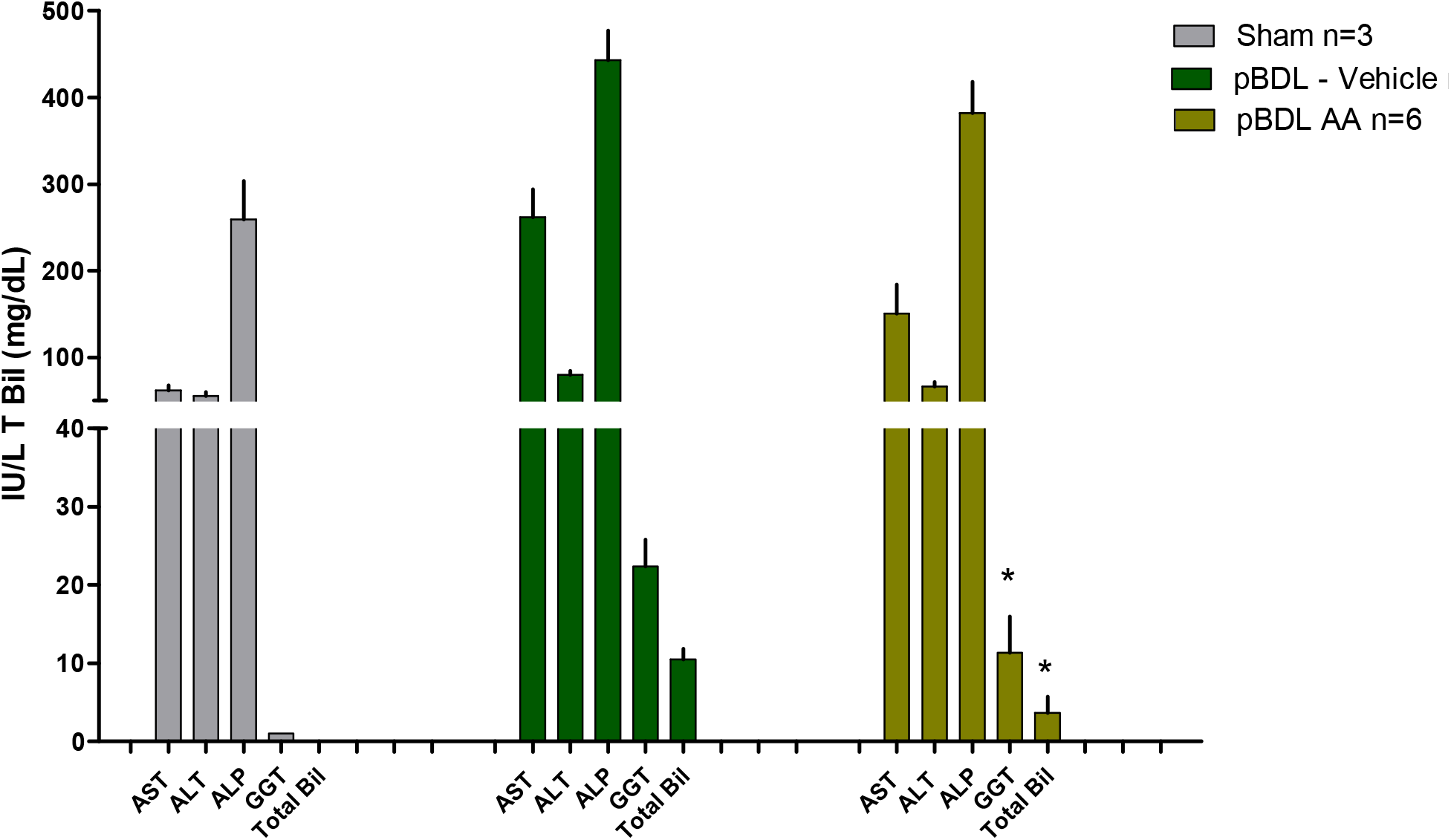
Day 14. Effect of AA on AST, ALT, ALP, GGT and total bilirubin levels in rats 14 days after pBDL surgery. Sham control data is provided to show effect of pBDL.

## 5. Conclusion

Hydrogen generated by AquaActive composition in rat stomach was determined to have a good efficacy for reducing key blood level biomarkers associated with liver injury suggesting the compound may provide novel option for the treatment of several liver diseases by decreasing of accumulation of toxins and reduction of levels of serum liver enzymes. We believe that the approach we are using is a simple and controlled method of hydrogen therapy. The main source of molecular hydrogen in the AA mixture is the metal particles of magnesium. Therefore, we believe that the use of similar mixtures with metallic magnesium can lead to similar therapeutic effects.

## Acknowledgements

The authors thank Dr. Christian S. Yorgure for reading the manuscript and helpful comments. This study was funded in part by Dibal, LLC (San Diego). The authors dedicate this article to the memory of Prof. Dr. Dusan Miljkovic, former CEO of Dibal, LLC, one of the pioneers of hydrogen therapy

## HIGHLIGHTS

- Simple and controlled method of hydrogen therapy for the liver injury
- Our acute model mimics many conditions of liver injury

